# HashSeq: A Simple, Scalable, and Conservative *De Novo* Variant Caller for 16S rRNA Gene Datasets

**DOI:** 10.1101/2021.01.29.428714

**Authors:** Farnaz Fouladi, Jacqueline B. Young, Anthony A. Fodor

## Abstract

16S rRNA gene sequencing is a common and cost-effective technique for characterization of microbial communities. Recent bioinformatics methods enable high-resolution detection of sequence variants of only one nucleotide difference. In this manuscript, we utilize a very fast HashMap-based approach to detect sequence variants in six publicly available 16S rRNA gene datasets. We then use the normal distribution combined with LOESS regression to estimate background error rates as a function of sequencing depth for individual clusters of sequences. This method is computationally efficient and produces inference that yields sets of variants that are conservative and well supported by reference databases. We argue that this approach to inference is fast, simple, scalable to large datasets, and provides a high-resolution set of sequence variants which are less likely to be the result of sequencing error.

## Introduction

Amplicon sequencing is a popular and cost-effective method for investigating microbial communities. A challenging step in using amplicon sequencing to identify members of microbial communities is to infer true sequences from artifacts. Sequence errors commonly occur both during PCR amplification and DNA sequencing. These errors include single nucleotide substitutions and gap errors due to mismatching bases and polymerase slippage, respectively^1^. For many years, standard practice was to lump sequences with 97% identity together into Operational Taxonomic Units (OTUs) in order to reduce noise and cluster closely related taxa^2–4^. However, recently developed bioinformatic tools attempt to infer true biological sequences at 100% identity by estimating the error profile and correcting point errors in sequences through denoising processes^1,5,6^. These pipelines rely on different assumptions and implement various statistical models. For example, DADA2 models error rate as a function of quality scores for each possible nucleotide transition and then these error rates are used in a Poisson-based model to infer true sequences from sequence errors^5^. Deblur compares sequence-to-sequence hamming distances to an upper-bound error model combined with a greedy algorithm^6^. Unoise2 uses two parameters that are pre-set values and are used for filtering low abundant sequencing and clustering of sequences based on their abundances^1^. All of these algorithms provide a higher resolution of taxonomic composition of a microbial community compared to the traditional OTU picking approach.

Despite the important progress that they represent, these algorithms all have some limitations. Deblur depends on construction of a multiple-sequence alignment which means that it does not scale to an entire dataset but works instead on each sample individually. This leads to the possibility of dependencies on the sequencing depth of each sample where variants might be called as real or artifactual differently in different samples depending on the properties of individual samples. Since Deblur sorts the abundance of sequences in each sample individually, it is also possible that the relative abundance of each variant within each sample can impact overall variant calling in complex ways. Deblur also has a number of free parameters and it is not immediately obvious how to optimize these parameters for new sequence datasets that might have different properties from the Illumina MiSeq and HiSeq training sets that were used for setting Deblur’s default values. Unoise2 is not freely available and also requires user-setting of parameters for which optimal values may not be entirely clear. As we will show, DADA2 can sometimes in practice yield larger numbers of sequence variants than can be considered biologically reasonable and often requires additional filtration of low abundance variants. Since DADA2 uses the Poisson distribution, it assumes that processes that control errors have similar rates for high and low abundance variants. These sorts of assumptions can be problematic in genomics. For example, in RNA-seq analysis it has long been understood that the relationship between mean and variance can have a relationship that is dependent on sequencing depth ^7^.

Here, we present HashSeq a very simple and fast algorithm for inferring sequence variants. We demonstrate that with enough sequence depth every possible unique one-mismatch variant for a sequence will be observed. We propose that the inference of true variants can therefore be determined relative to this background probability of observing one-mismatch variants, which can be approximated with a two-parameter normal distribution. We applied this method to six publicly available datasets and show that this simple approach is fast and scales well even to large datasets. Our approach provides a conservative set of variant calls that is well supported by a reference database and behave almost identically to DADA2 calls in supervised classification.

## Results

### With sufficient sequencing depth, all one-mismatch “child” variants for a “parent” sequence are likely to be observed and this is well modeled by a simple Poisson process

We used a HashMap data structure, which identifies every unique sequence in linear time proportion to the total number of sequences, to identify all sequence variants in six publicly available Illumina datasets. Sequences from these projects were obtained from three fecal microbiota (China, Autism, and RYGB datasets), one vaginal microbiota, and one soil microbiota dataset as well as one microbial mock community (MMC, see methods). This method of sequence variant detection is very fast (less than 1 hour even for the largest dataset with 416,450,026 sequences), but it results in a large number of sequence variants ranging from 6,166 for the smallest dataset (mock community) to 814,494 for the largest dataset (Vaginal dataset). The majority of these variants are presumably sequencing errors or other artifacts. In order to detect sequence errors, we clustered sequences that had only one nucleotide difference (Figure 1). Under this approach, sequence variants were sorted according to their abundances. Starting with the most abundant sequence variant, considered as the first “parent sequence”, clusters were formed by adding all the one-mismatch variants to each cluster (one-mismatch children). This resulted in 2,002 clusters of parents plus children (when present) for the smallest dataset (MMC dataset) and 387,903 clusters for the largest dataset (Vaginal dataset). We assessed the relationship between the abundance of parents and the number of one-mismatch children present in each cluster. Figure 2 shows the fraction of all possible one-mismatch children as a function of the abundance of each parent sequence for six datasets. When the abundance of a parent sequence is high enough, almost all possible unique one-mismatch children for that parent sequence can be observed (Figure 2). For example, the read lengths for both the China and Vaginal datasets are 250 bp, therefore, there are 750 possible one-nucleotide differences for a parent sequence in these datasets. For these datasets, the most abundant parent sequences have more than 10,000 reads and almost all of the possible child variants were observed (the rightmost points in Figure 2). For the least abundant parent sequences (the leftmost points in Figure 2), almost no one-mismatch variants were observed.

**Figure 1.**
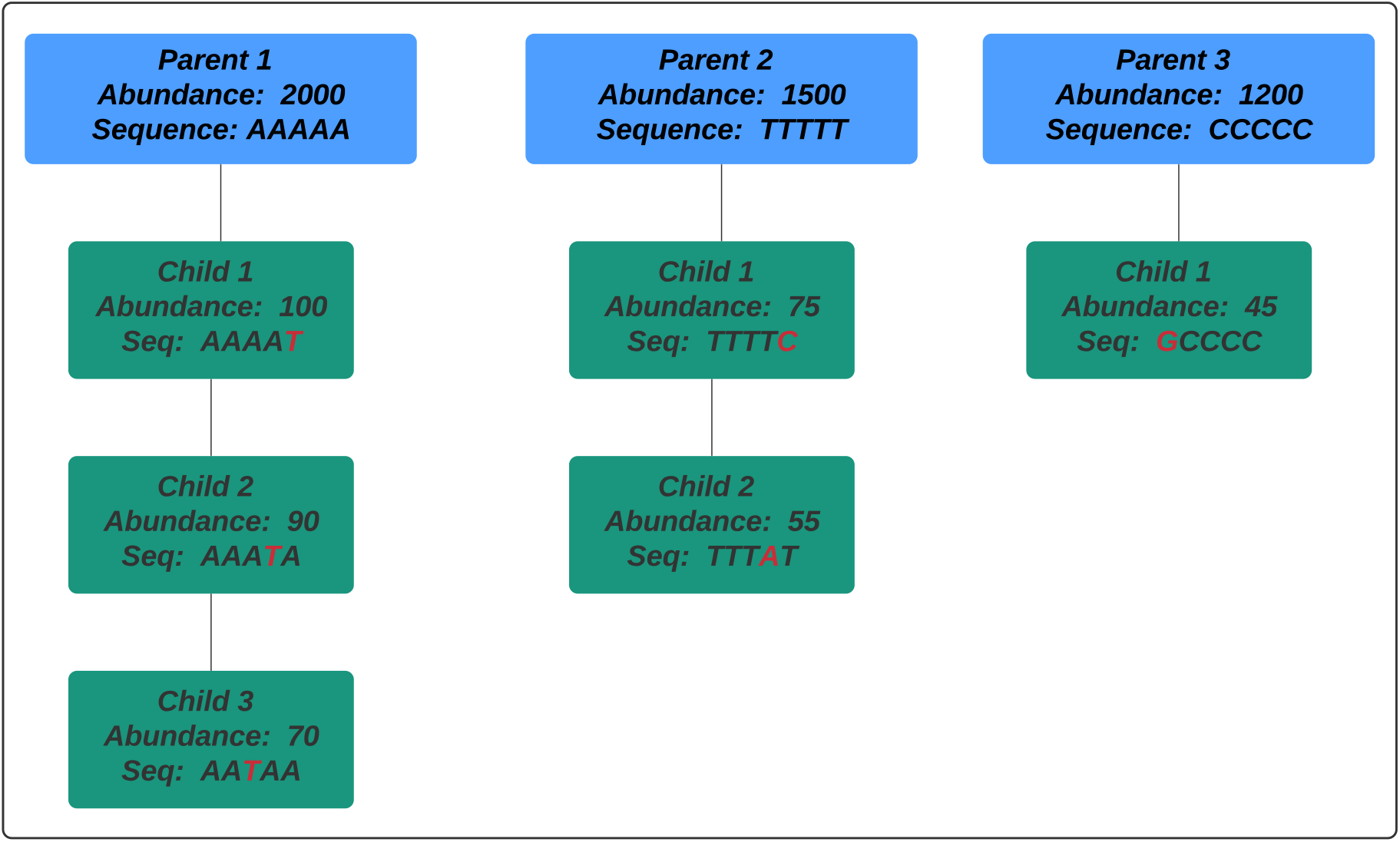
Cluster formation of parents and their one-mismatch children in the HashSeq algorithm. In this clustering strategy, sequence variants are sorted according to their abundances. Starting with the most abundant sequence variant, considered as the first “parent sequence”, clusters are formed by adding all the one-mismatch variants (one-mismatch children) to each cluster.

**Figure 2.**
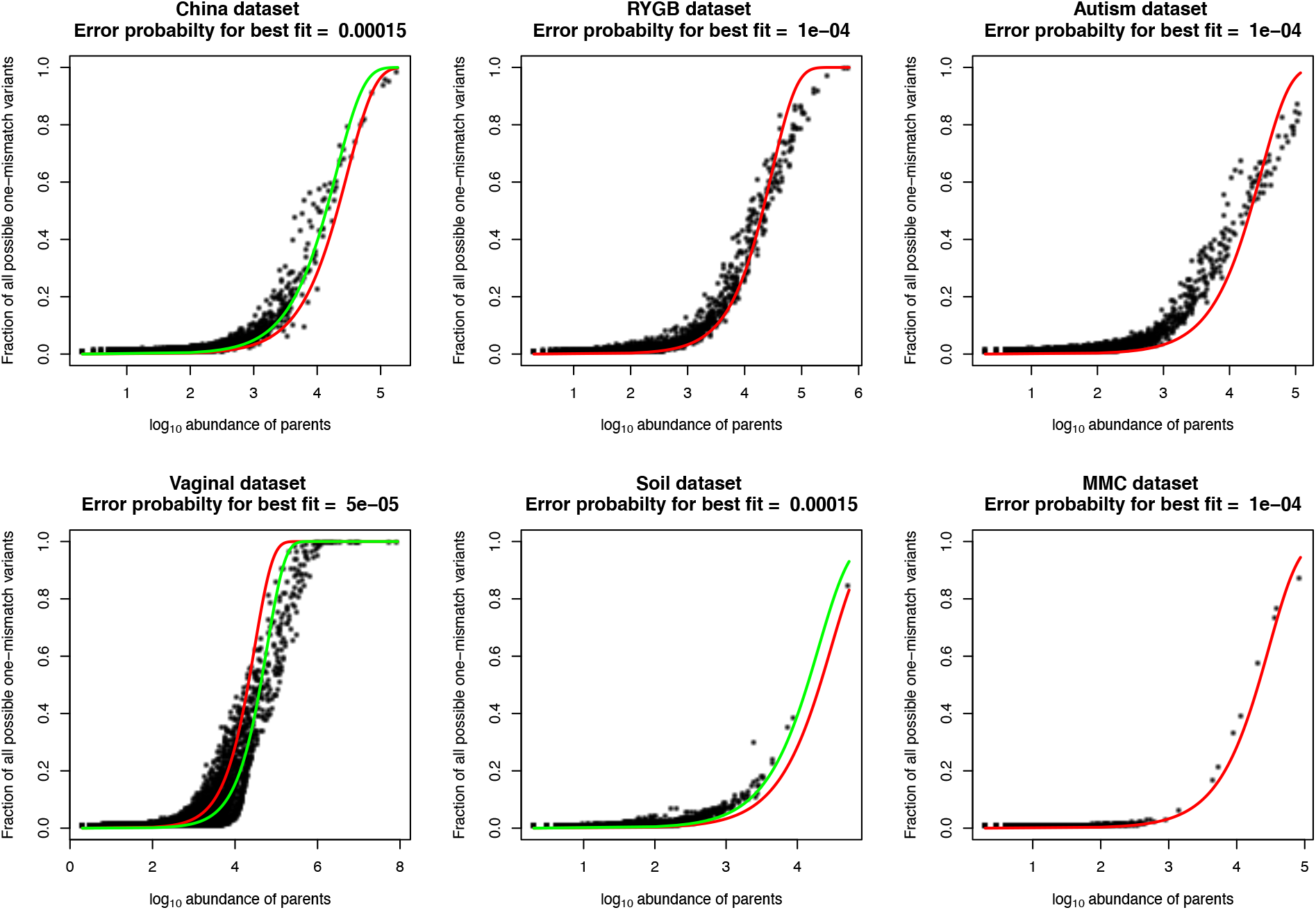
The presence or absence of unique one-mismatch variants can be surprisingly well modeled with a simple one-parameter Poisson distribution with an almost constant error-rate across six independent 16S rRNA gene Illumina datasets. Plots show the relationship between the abundance of parent sequences on the log 10 scale and the fraction of all possible unique one-mismatch variants for the parent sequences. These data are well modeled by a simple one-parameter Poisson distribution. The red line corresponds to an error rate of *p* = 10^-4^. The China, Vaginal, and Soil datasets were best modeled using slightly different error rates for each dataset (green lines, China and Soil *p* = 1.5 *10^-4^, Soil *p* = 5*10^-5^).

Interestingly, these data were surprisingly well fit with a simple Poisson distribution with a single parameter across all datasets. The single parameter is the probability (*p*) that a single nucleotide will be different between two sequence variants (see methods). Even though this model does not contain any information about different error rates for different nucleotides, or any information about the biology of any of these diverse ecosystems, all the datasets were reasonably well fit with a parameter of *p*=10^-4^ (Figure 2, red curves) although for some datasets a slightly better fit could be obtained with a slightly different value for *p* (Figure 2, green curves). The consistency of this fit across datasets is perhaps surprising given that not all the datasets used the same primers for PCR amplification as well as the wide biological variability of these samples extending from the vaginal to the gut and soil microbiome. This analysis suggests that a common baseline error-rate exists across multiple Illumina datasets and that the probability of seeing a one mismatch variant is well-modeled as a simple function of the abundance of the parent sequence. Our results demonstrate that with enough sequencing depth, every possible one-mismatch child is likely to be observed for all variants and in the absence of any other information, it is possible to predict the likelihood of seeing a unique child variant given only the abundance of the parent.

### The background Poisson distribution underestimates the true abundance of one-mismatch “child” variants, while a normal distribution-based model provides a better fit

Since we have demonstrated that the number of one-mismatch variants accumulate as a simple function of sequencing depth, the challenge for all algorithms in finding sequence variants is to discriminate true variants from the many stochastically produced artifactual variants. One possible approach to this problem might be to use the estimated error rate derived from the presence or absence of one-mismatch variants (as described in the previous section) to predict the background abundance of sequence errors and only consider “true” variants if the abundance of sequence variants is significantly enhanced over the expected background noise. However, when we tried to use this background error rate as a threshold for determining true variants from artifacts using the Poisson test (see methods), we rejected the null hypothesis that the sequence variant was due to random sequencing error for more than 83% of one-mismatch children even after correcting for multiple hypothesis testing (Supplementary Figure 1). This suggests that the distribution of children abundance does not follow the Poisson distribution. Indeed, the Poisson distribution assumes that the mean equals the variance, and clearly this assumption does not hold as the variance of children abundance shows clear over-dispersion, that it is larger than the mean of children abundance for most parent sequence variants across datasets (Figure 3). As a result, the Poisson distribution underestimates the true variance and is therefore anti-conservative and call nearly all one-mismatch variants as true variants.

**Figure 3.**
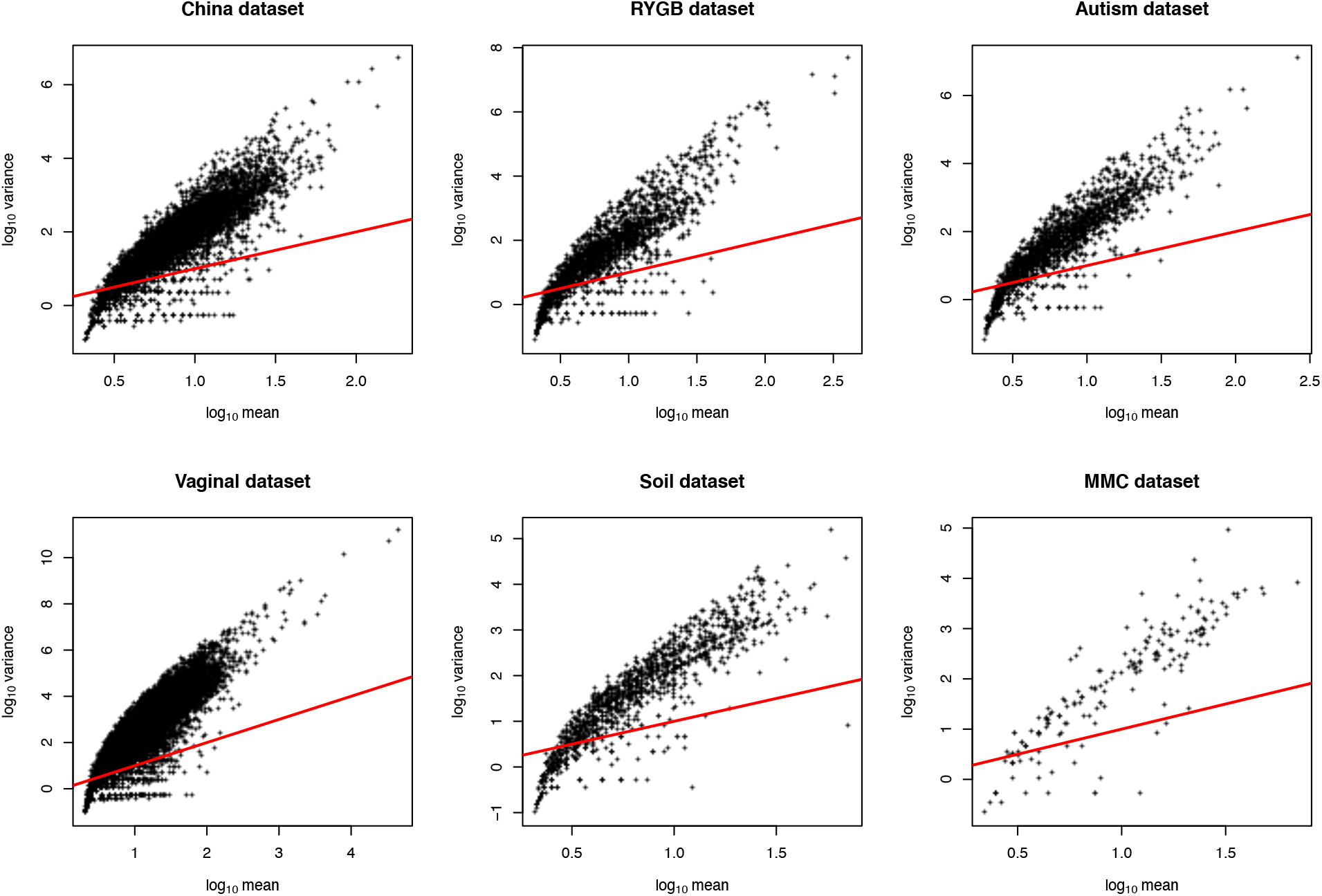
The variance of the one-mismatch children abundance in each cluster is not equal to their mean abundance. Plots show the relationship between the variance and mean abundance of one-mismatch children from each parent cluster across six different 16S rRNA gene datasets. The red line represents the Poisson assumption of equal mean and variance.

As we observed that the Poisson distribution appears to be extremely anti-conservative, we next examined whether the distribution of one-mismatch children could be better explained by a normal distribution since it is more flexible in terms of the relationship between the mean and variance compared to the Poisson distribution. For this, the abundances of children were log_10-_ transformed and the distributions of log_10_ – transformed abundances of children were plotted for each parent sequence. The histograms of children abundances (shown for the most abundant parent for each dataset in Figure 4) suggest that the distribution of one-mismatch children approximates a normal distribution. Interestingly, we observed that the mean abundance of children for each cluster can be well fit by a Locally Estimated Scatterplot Smoothing (LOESS) function of the parent abundance, especially for high abundance parents (>1,000 reads) across all the six datasets (Figure 5). The smooth relationship between mean and standard deviation and sequencing depth across the six datasets suggests that there is a general error rate across all variants that is dependent on sequencing depth but not dependent on the biology of each particular parent sequence. This further suggests that the LOESS fit may represent a good model for general inference. However, when parents have a lower abundance, generally below 1,000 sequences across all samples, a smaller number of one-mismatch variants are present (Figure 1) and therefore variance in the mean abundance of children significantly increases (Figure 5) presumably due to sparsity effects.

**Figure 4.**
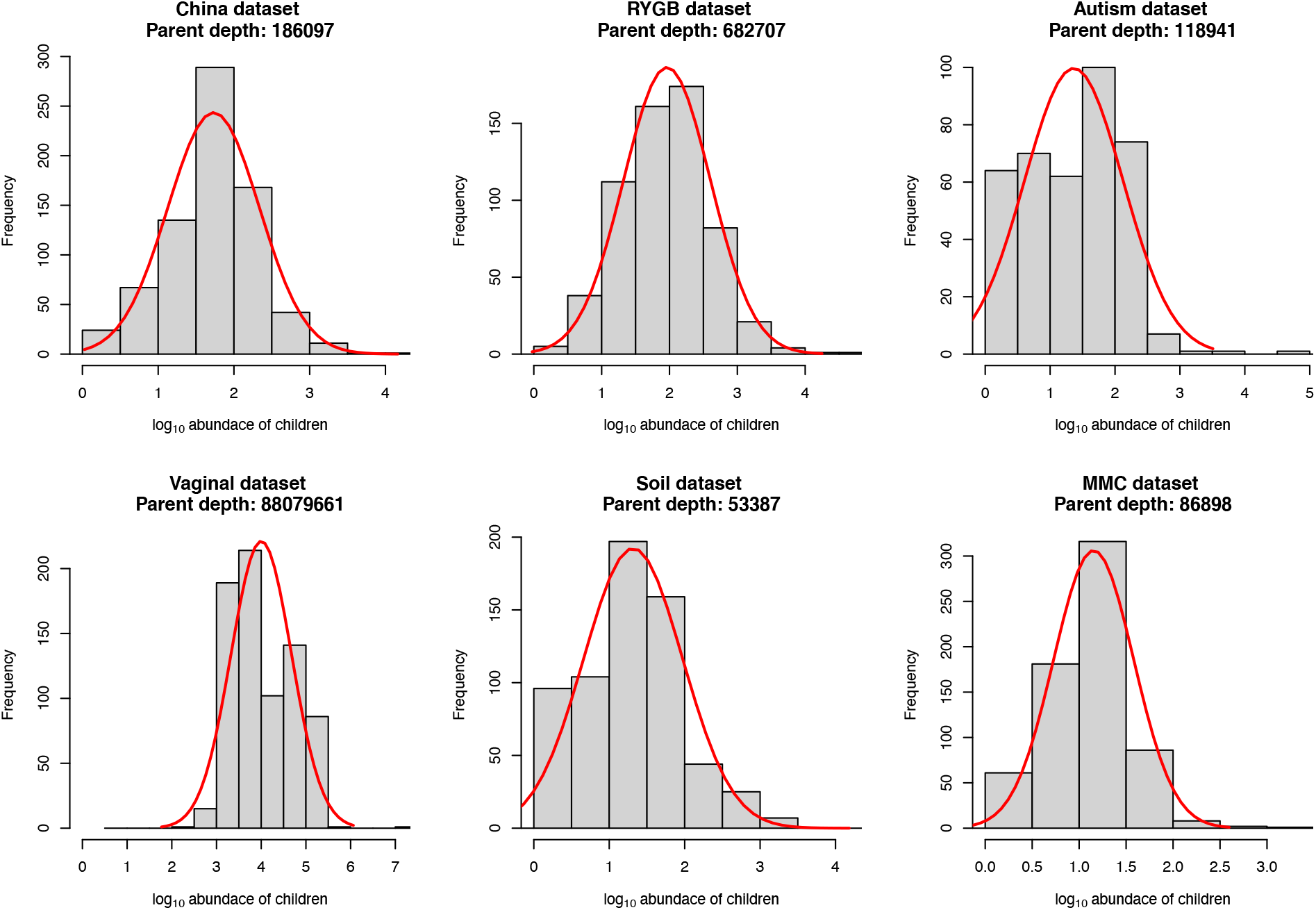
The abundance of one-mismatch children within a cluster is approximately normal on a log_10_ scale. Histograms showing the distribution of abundance of one-mismatch children for the most abundant parent on a log_10_ scale across the six different 16S rRNA gene datasets.

**Figure 5.**
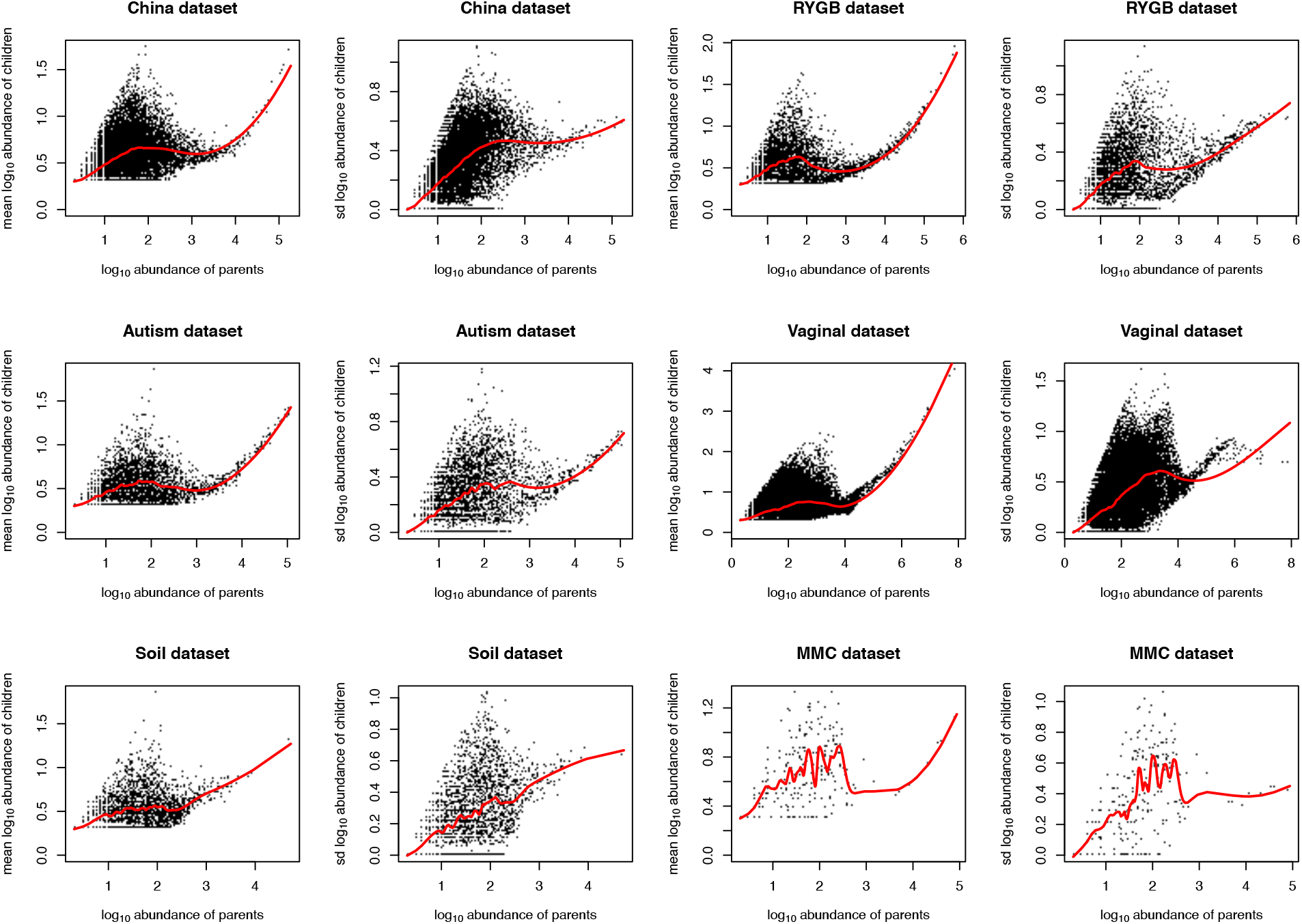
Mean and standard deviation of one-mismatch children in each cluster is a smooth function of their parent abundance on a log_10_ scale for the most abundant parent sequences. Plots show the relationship between the mean and standard deviation of one-mismatch children abundance in each cluster and their parent abundance on a log_10_ scale. The mean and standard deviation of abundance of one-mismatch children for each cluster can be well fit by a smooth LOESS function of the parent abundance especially for high abundance parents (>1,000 reads) across six different 16S rRNA gene datasets (red line).

### Normal-based Inference of one-mismatch “children” is fast, conservative and produces comparable results to DADA2 in supervised classification analyses

The above results suggest that we can assume that the background distribution of children variants is reasonably normally distributed and is well fit for sequences with abundance >~1,000 reads by a simple localized regression (or LOESS). In this section, we explore an inference scheme in which the background mean and standard deviation are the higher of the mean and the standard deviation found for each parent (black dots in Figure 5) or the LOESS regression of the mean and standard deviation (red lines In Figure 5). In this scheme, we use these estimates of mean and standard deviation as our background null hypothesis that the abundance of the one-mismatch child variant is a sequence error and can therefore be explained by the background level of sequencing error of the parent. From these background mean and standard deviation, we generate a one-sided p-value (using “pnorm” in R) for rejecting the null hypothesis. A small p-value for this null hypothesis indicates that a child variant has an abundance level above this expected background for its parent (see methods).

When using a 5% false discovery rate, this method results in a considerably lower number of sequence variants compared to DADA2 with default parameters for the non-mock biological datasets (Table 1) and compared to the inference test based on the Poisson distribution described above (Supplementary Figure 1). When we mapped the inferred sequence variants with BLAST to the SILVA132 dataset, the great majority of sequence variants had a high degree of identity (>99%) to the SILVA database (Table1 and Figure 6) suggesting that many of the variants that we detected had been previously observed. This supports an assertion that these variants are not sequencing error. Interestingly, although HashSeq calls more variants in the MMC dataset compared to DADA2, the parent sequences include the eight bacterial taxa that are present in the mock (*Listeria monocytogenes, Pseudomonas aeruginosa, Bacillus subtilis, Escherichia coli, Salmonella enterica, Lactobacillus fermentum, Enterococcus faecalis*, and *Staphylococcus aureus*), which further confirms that our clustering strategy is able to find major taxa in a dataset. Overall, these results suggest that our normal distribution-based inference approach is often more conservative compared to DADA2 and less prone to infer superius variants as “true” sequences. However, because we do not look for variants in low-abundance regions where the LOESS regression does not show a consistent relationship to sequencing depth, our algorithm is less sensitive to low abundant true sequences.

**Table 1.** Results of mapping of sequence variants inferred by HashSeq and DADA2 to the SILVA132 database using BLAST. The table shows the number of sequence variants inferred by DADA2 and HashSeq as well as the percent Identity of the sequence variants to the SILVA132 database.

| Dataset | Pipeline | #SVs | Parents |  |  |  |  | Children |  |  |  |
| --- | --- | --- | --- | --- | --- | --- | --- | --- | --- | --- | --- |
|  |  |  | Parents-Children | Identity=100 % | 100 % > identity ≥ 99 % | 99 % > identity ≥ 97 % | 97 % > identity | Identity=100 % | 100 % > identity ≥ 99 % | 99 % > identity ≥ 97 % | 97 % > identity |
| China | HashSeq | 451 | 255-196 | 194 | 34 | 21 | 6 | 67 | 102 | 23 | 4 |
|  | DADA2 | 4974 | --- | 1050 | 540 | 400 | 2984 | --- | --- | --- | --- |
| RYGB | HashSeq | 718 | 402-316 | 373 | 21 | 5 | 3 | 147 | 169 | 0 | 0 |
|  | DADA2 | 3385 | --- | 1648 | 966 | 191 | 580 | --- | --- | --- | --- |
| Autism | HashSeq | 422 | 229-193 | 214 | 8 | 6 | 1 | 82 | 101 | 9 | 1 |
|  | DADA2 | 2398 | --- | 1295 | 493 | 230 | 385 | --- | --- | --- | --- |
| Vaginal | HashSeq | 1305 | 530-775 | 222 | 191 | 72 | 45 | 68 | 400 | 195 | 112 |
|  | DADA2 | 12106 | --- | 3284 | 4335 | 1562 | 2925 | --- | --- | --- | --- |
| Soil | HashSeq | 49 | 39-10 | 37 | 1 | 1 | 0 | 9 | 1 | 0 | 0 |
|  | DADA2 | 5761 | --- | 1256 | 1387 | 1466 | 1652 | --- | --- | --- | --- |
| MMC | HashSeq | 35 | 9-26 | 9 | 0 | 0 | 0 | 5 | 21 | 0 | 0 |
|  | DADA2 | 26 | --- | 14 | 8 | 1 | 3 | --- | --- | --- | --- |

**Figure 6.**
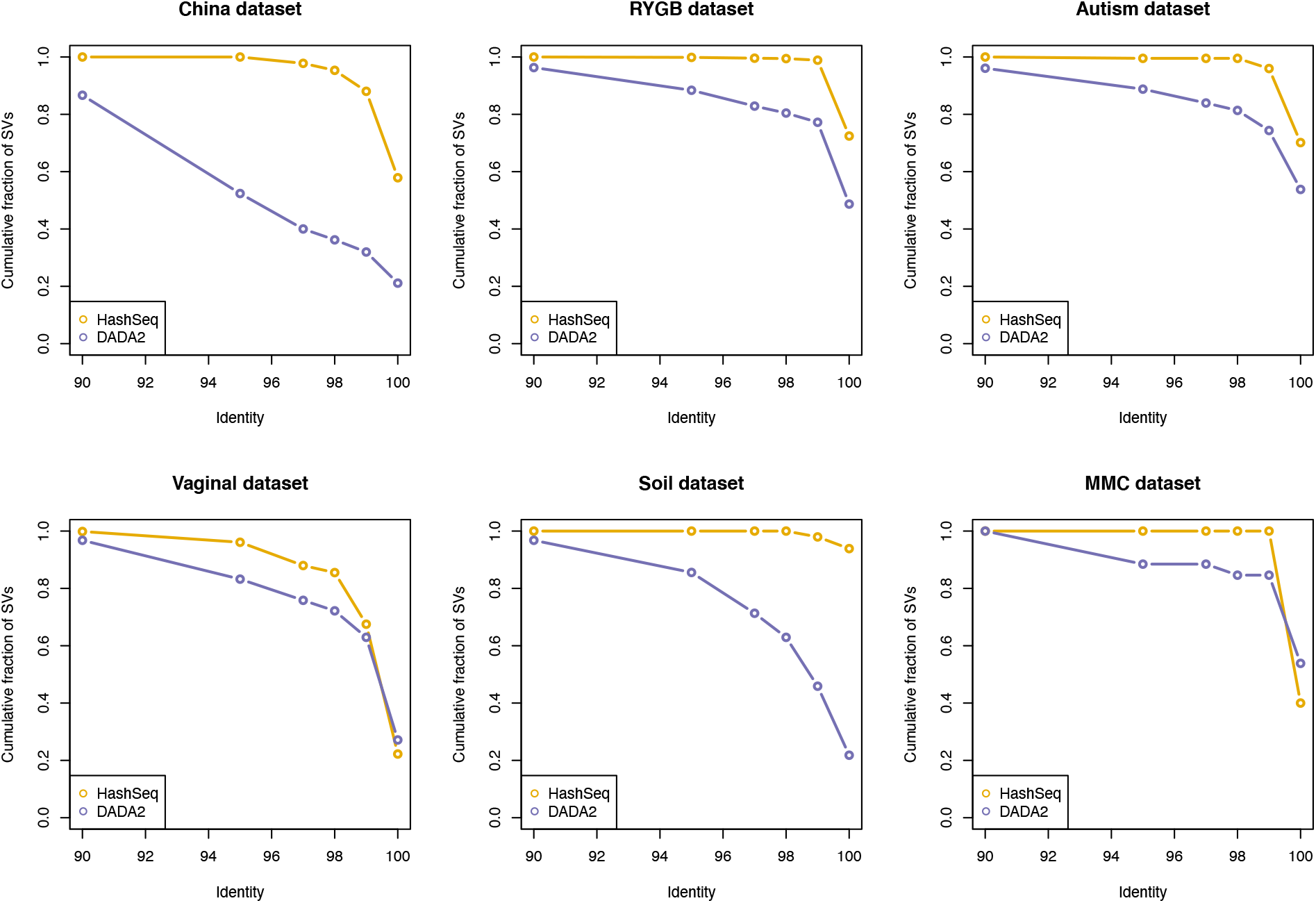
Sequence variants generated by the HashSeq pipeline have a high degree of identity to the SILVA132 database. For each dataset, the cumulative fraction of inferred sequence variants for a range of 90-100% identity to the SILVA132 database is plotted for both the HashSeq and DADA2 pipelines.

Next, for each dataset we performed a Random Forest classification to study the association between the sequence variants with metadata variables of interest in the five publicly available non-mock datasets. Compared to DADA2, our approach performs nearly identically in terms of association studies between the gut microbiota and biological variables (Figure 7).

**Figure 7.**
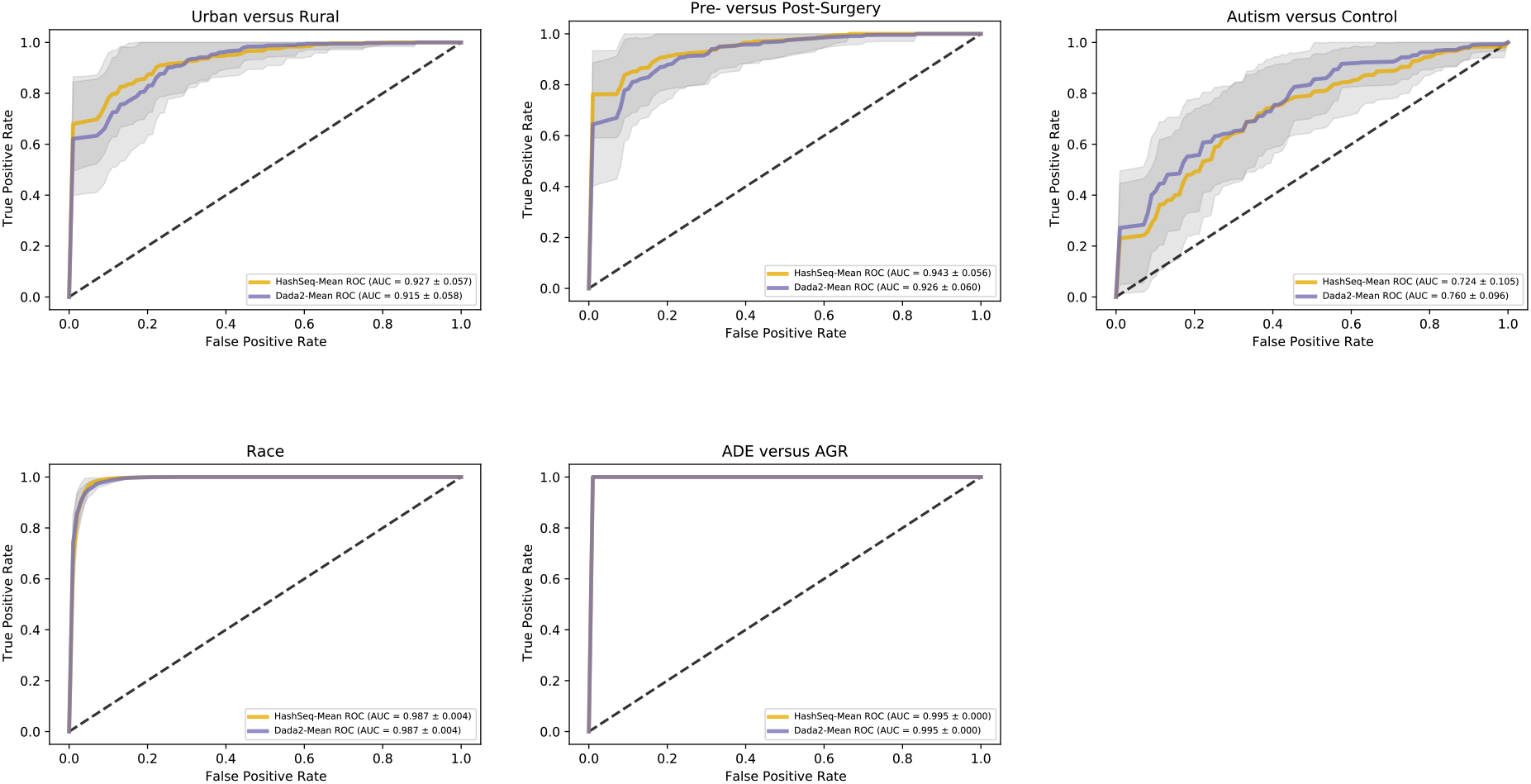
HashSeq performs almost identically to DADA2 in terms of association studies between the gut microbiota and biological variables. Random forest classification was used to study the association between the sequence variants with metadata variable of interest for five different 16S rRNA datasets. The area under the curves (AUC) of the ROC curves were essentially superimposable between our inference-based approach and DADA2. For the China dataset, we examined if the gut microbiota can predict rural versus urban samples. For the RYGB dataset, we tested if the gut microbiota can predict pre-surgical versus post-surgical samples. For the Vaginal dataset, we studied the association between the microbiota and ethnicity (black women versus white women). For the Autism dataset, we examined if the gut microbiota can predict children with Autism versus the control group. For the Soil dataset, the association between the microbiota and two types of Soil, Amazon Dark Earth (ADE) and agricultural Soil (AGR) was examined.

Finally, we compared run-time and memory usage between our pipeline and DADA2. On average across datasets, our pipeline is 43 times faster than DADA2 and the memory usage is 3 times less than DADA2 (Figure 8). For the Vaginal dataset, the largest dataset, HashSeq was 6.5 times faster than DADA2; however, it was comparable to DADA2 in terms of memory usage.

**Figure 8.**
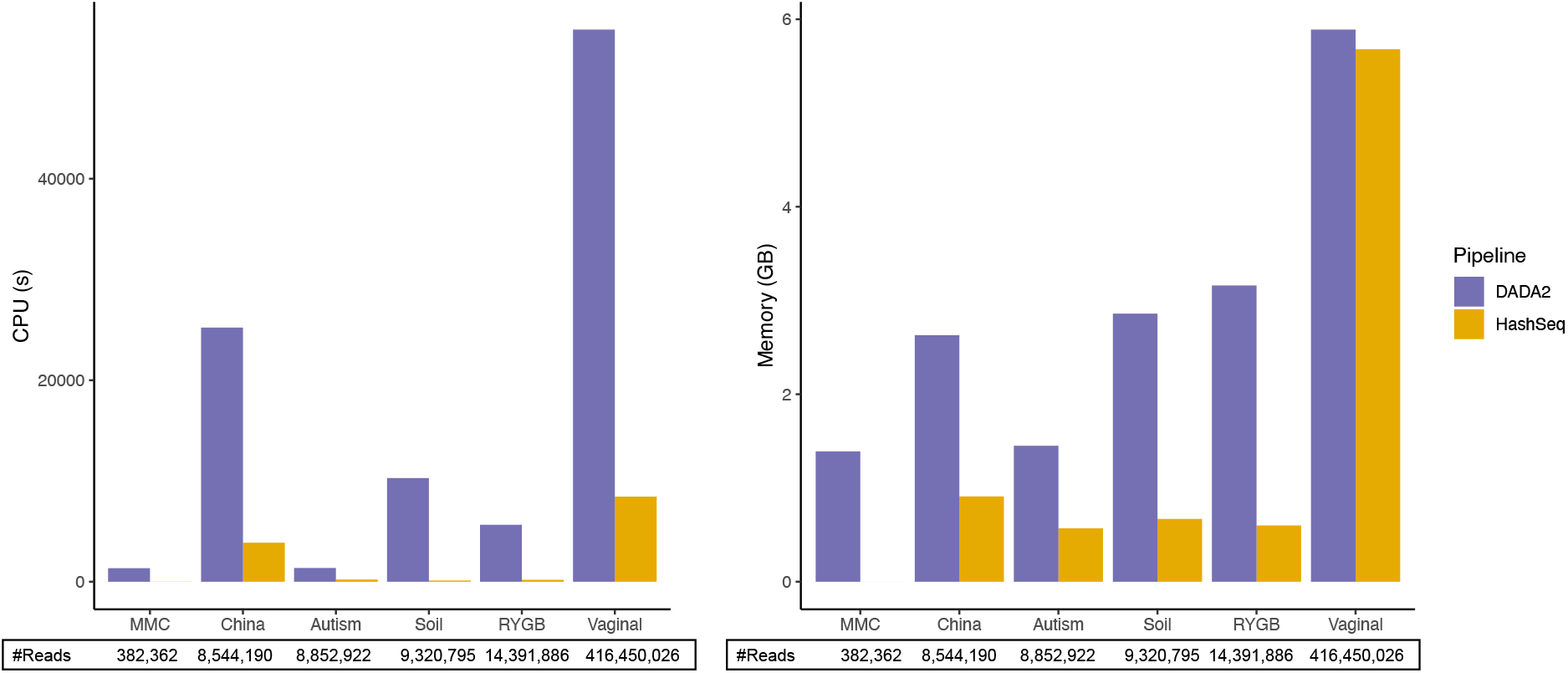
HashSeq is faster and more efficient in memory usage compared to DADA2. Run-time and memory usage by DADA2 and HashSeq are plotted across for each 16S rRNA gene datasets.

## Discussion

In this paper, we utilized a very simple HashMap based algorithm to detect all sequence variants in a dataset. This resulted unsurprisingly in a large number of one-mismatch sequence variants. We assume that nearly all these spurious sequences are caused by sequencing error. Our paper provided two lines of evidence to support this assertion. First, the number of distinct one-mismatch children for each parent sequence can be well modeled by a simple Poisson process, suggesting that when sequence depth is high enough every possible one-mismatch variant of a parent sequence can be observed. This seems unlikely to be explained by biological variance. A second line of support for the assertion that most variants are related to sequencing error is the excellent fit to a smooth LOESS curve with sequencing depth over 1,000 sequences (Figure 5), suggesting that sequencing depth and not the biology of a particular cluster control the abundances of observed variants.

Given a postulate that nearly all sequence variants are the result of error, a natural approach is to use the background error rate for inference to detect the relatively rare occurrence of a variant that cannot be explained by background sequencing error. This approach of using a background rate to generate p-values for an event that is reasonably uncommon has long been an approach to inference in genomics^8^. A natural question is how to parametrize the expected background rate. Since we know that there is a dependency on sequencing depth, the simplest approach would be a Poisson based model in which the mean equals the variance. However, the Poisson model failed to control the distribution of one-mismatch children in each cluster presumably because the Poisson assumption of equal mean and variance is not met. In a similar way, previous studies have shown that when the Poisson distribution is used to test for differential gene expressional in RNA-seq datasets, the Poisson estimated variance is smaller than the observed variance in real data, resulting in increased false discoveries^7,9^. Therefore, over-dispersion (where variance is higher than mean) is a general feature of sequence count genomic datasets, including sequence variants, and this problem causes inference based on the Poisson distribution to fail.

We therefore argued that inference of a true variant should utilize a model in which the variance was not constrained to equal the mean. Previous algorithms designed for RNA-seq datasets, such as DESeq and EdgeR, model count data with a negative binomial distribution which assumes the variance is greater than the mean^7,10^. We preferred the normal distribution over the negative binomial distribution to model the background error for two reasons. First, the negative binomial distribution is not defined when the variance is less than mean, and although for the majority of sequence variants the variance is greater than the mean, there are still a large number of children sequences that have a mean greater than the variance (Figure 3). Second, the negative binomial as a count model does not work on log-transformed data, which contains noninteger values. The normal distribution instead gives us more flexibility in terms of the relationship between the mean and variance as well as more naturally allowing for the transformation of count data. Regardless of the limitations of the negative binomial distribution, at high sequencing depth the negative binomial distribution is well approximated by the normal distribution, further justifying the use of the normal distribution.

In order to use a normal distribution-based model to infer true sequences from the background noise, we used the mean and standard deviation predicted by a localized regression fit between mean and standard deviation and parent sequences (Figure 5). In order to be as conservative as possible, we chose the mean and standard deviation for our inference test to be the higher of the mean and the standard deviation found for each parent directly or predicted by the LOESS regression. This conservative approach detected sequence variants that had a good match to existing variants in the SILVA database, suggesting that many of the variants that we detected had been previously observed and therefore are unlikely to be sequencing error. This provides further confirmation of the conservative nature of our method.

Our normal-distribution-based algorithm for detection of sequence variants which we here call HashSeq has a number of advantages. First, it is very fast and can detect sequence variants in less than three hours on a single CPU even for a very large dataset. It can run all sequences in a dataset together and does not require running sequences from each sample independently. This eliminates any potential problems in which the characteristics of individual samples impact overall variant calling in potentially complex ways. Second, our algorithm compared to the popular algorithm DADA2 is fairly conservative and calls a fewer number of sequence variants as true. The conservative nature of our method potentially increases the power of a study to detect a signal since fewer number of spurious variants are reported and therefore there will be a fewer number of hypothesis to be corrected for in downstream analyses using FDR multiple hypothesis correction. By determining where the smooth relationship between parent abundance and mean of children abundance breaks down, our algorithm offers a natural way to set a threshold for removing low abundance variants.

This was set at a parent abundance of 1,000 reads for all of our datasets except the largest one, where it was set to 10,000. Setting a low abundance threshold in this way is an appealing alternative to removing taxa based on arbitrary thresholds of rarity or prevalence. In addition, because our algorithm provides explicit p-values for a null hypothesis that a child sequence was derived from sequencing error of a parent sequence, our results may be easier to interpret than algorithms that do not assign a score to variants or assign scores based on arbitrary scales.

Finally, our algorithm is simple and is based on a two-parameter model. By contrast, DADA2 assumes that each nucleotide transition has its own parameter which is calculated from the transition probabilities and quality scores. DADA2 assumes that the parameters obtained from quality scores are independent of sequencing depth while our model explicitly considers background mean and variance as a function of sequencing depth. Despite these differences in parametrization, using our variants or DADA 2 variants produces essential identical power for machine learning based supervised classification.

Our algorithm has some limitations to be noted. First, our algorithm is not sensitive to detect true low-abundance sequences. Therefore, we recommend using more sensitive algorithms, such as DADA2, to detect low-abundance sequence variants. Another limitation is that our algorithm does not consider two- or more mismatches. However, we believe that one-mismatch errors are more likely to happen compared to two- or more mismatches and therefore the abundances of two- or more mismatches will reliably fall below the detection limit of the algorithm (usually <1,000 reads). This assertion is supported by simulation results (supplementary Figure 2) which suggest that variants with more than one mismatch error occur very infrequently in short-reads. This assumption of the rarity of multiple-mismatch sequences, however, may not be appropriate for long-read technologies such as PacBio, and this is another potential limitation of our method. Finally, our algorithm does not explicitly model insertions or deletions (indels) and will treat indel events as a separate parent sequence. Users who need to capture indel variation in relationship to a parent might consider use of Deblur or other methods that incorporate multiple sequence alignments.

In summary, we described HashSeq, a very simple and fast algorithm to infer true variants from background sequencing error. This algorithm can be easily used for small or large 16S rRNA gene datasets generated from a diverse range of ecosystems. Source code is freely available at https://github.com/FarnazFouladi/HashSeq as an R package.

## Methods

### Publicly available datasets

Six datasets were included in this study: one publicly available microbial mock community (MMC) consisting of three samples (PRJEB24409) and five publicly available 16S rRNA gene datasets, including three human gut microbiota datasets to which we refer as “China” (PRJNA349463, n=80), “Autism” (PRJNA533120, n=81), and “Roux-en-Y Gastric Bypass (RYGB)” (SRP113514, n=71), one Vaginal microbiota dataset (SRP115697, n=2367), and one Soil microbiota dataset (PRJEB14409, n=40)^11–16^. For all datasets except for the Soil dataset, forward and reverse reads were merged using the PEAR software^17^, and the paired reads were then trimmed to a constant length and shorter reads were discarded (250 nucleotides for China, Vaginal, and MMC, 200 nucleotides for RYGB dataset, and 151 nucleotides for Autism dataset). For the Soil dataset, only forward reads were used, due to concerns about sequence quality for the reverse reads, and the reads were trimmed to 250 nucleotides. For the Soil, RYGB, and MMC datasets primers were present in the public sequences and were removed by our pipeline. Information regarding primers and the variable region of 16S rRNA gene that were sequenced can be found in the Supplementary Table1. For all datasets, singletons in each sample as well as sequences with N’s were removed.

### Cluster of sequences composed of a parent and one-mismatch children

We used a HashMap, a simple data structure, to detect all 16S rRNA gene sequence variants, excluding sequence variants with only one read in a sample (singletons). In our method, sequence variants are sorted according to their abundance. Starting with the most abundant sequence variant (considered as a “parent” sequence), all possible one-mismatch sequence variants in the dataset are identified and considered as the one-mismatch “children” for that parent sequence. Similar searches for parents and children sequences are performed for the remaining sequences until all sequences are assigned as a parent sequence or as a one-mismatch child sequence, resulting in the formation of numerous clusters of sequences that are composed of one parent and one-mismatch children (Figure 1).

### Poisson model of frequency of one-match children variants

In order to estimate the rate of observing a one-mismatch sequence variant, we fit our data to a very simple model based on the Poisson distribution. This model has one free parameter which is the probability (*p*) of a single-nucleotide sequencing error. In this approach, we treat each nucleotide within a set of parent and children sequence variants independently. Given the background error rate of *p* and a parent sequencing depth of P_i_ (a parent belonging to the cluster i), the probability of seeing at least one error for a given nucleotide is given by:

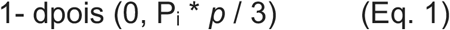

We divide *p* by 3 in the above equation because there are 3 possible distinct nucleotides that can be tabulated as an error (for example ‘A’ can be erroneously observed as either ‘C’, ‘G’ or ‘T). If *p* is close to 1, we would expect to always see all three possible variants of the nucleotide and if *p* is close to zero, we would expect to never see a mutation at that position in the sequence. We will argue that we can fit Eq. 1 to our datasets by considering the fraction of all unique one-mismatch children observed for a parent sequence divided by the number of all possible one-mismatch children as shown in Figure 1 as described in the results. If *p* is high, then we would expect to see most of the all possible one-mismatch variants of a parent sequence and if *p* is low, we would expect to see few.

This model makes a number of simplifying assumptions. A key assumption is that the error rate *p* can be estimated independently for each nucleotide and that not considering sequences with more than one nucleotide difference between parent and children does not bias our error rate estimate. We think this is a reasonable assumption as simulating a polymerase with the same error rate as the Poisson equation above and examining the resulting distribution of one-mismatch children observed from among all resulting sequences yields an essentially identical distribution as the Poisson equation above (the simulation code is available here: https://github.com/afodor/metagenomicsTools/blob/master/src/binomFit/HowManyVariants.java and Supplementary Figure 2). This concordance occurs because the overall error rate is low enough that sequences with more than one mismatch occur infrequently and can therefore be ignored without altering our baseline error rate estimate. For example, for a 250 base-pair sequence length with a *p =* 0.00015 errorrate, sequences with more than one mismatch are seen only in about 1 of 1,600 sequences in our simulation code. Obviously, this assumption of independence of sequence variants that allows us to ignore sequences with multiple mismatches becomes more problematic for read lengths greater than the 250 base-pair that we examine here and for overall higher error-rates.

In addition, in order to see whether the estimated error rate derived from the presence or absence of one-mismatch variants using the above model can be used to predict the background abundance of sequence errors and therefore to infer true variants whose abundances are above the background noise, we used a Poisson test using “poisson.test” in R with the following parameters:

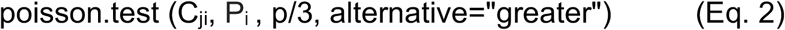

where C_ji_ is the abundance of a j^th^ child sequence of the cluster i, P_i_ is the abundance of a parent sequence of the cluster i and *p* is the estimated error rate from the Poisson model above (Eq. 1). P-values generated by the Poisson test (Supplementary Figure 1) were adjusted for multiple hypothesis testing using the Benjamini-Hochberg Procedure.

### Normal distribution of the background noise

As equation 2 based on the Poisson distribution underestimates the abundances of one-mismatch children (see result section), we further examined if the abundances of one-mismatch children can be better fit by the normal distribution. For this purpose, abundances of sequence variants were log_10_ transformed and their histograms were plotted for each parent sequence. Next, the mean and standard deviation for each parent were calculated. The relationship between the mean abundance of children and the parent sequences as well as the standard deviation abundance of children and the parent sequences were fitted to a local regression or Locally Estimated Scatterplot Smoothing (LOESS). We will show that for parent sequences with depths above > 1,000 reads, the LOESS regression is a reasonable fit for most datasets, however, for parents with depths below 1,000 reads, variance of the means and standard deviations are increased due to the sparsity of one-mismatch children, and therefore the LOESS regression does not fit as well (Figure 5). Therefore, as a default, sequence variants with total abundance less than 1,000 across all samples are filtered (i.e., removed) in our pipeline. This threshold of 1,000 can be changed by users based on their data. For example, for the Vaginal dataset we increased the threshold to 10,000 as the sequencing depth is significantly higher for this dataset compared to other datasets and the LOESS regression is a good fit when sequences have depths higher than 10,000 reads (see Figure 5).

The means and standard deviations estimated from the LOESS regression were assumed to be the background noise and therefore any variant above this background noise would be called a true sequence variant. Based on this assumption, for each child variant, a one-sided p-value was generated using the “pnom” function in R and the following formula:

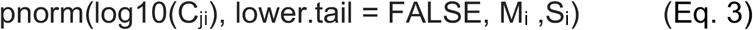

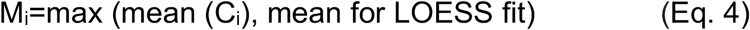

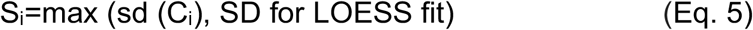

where C_ji_ is the abundance of j^th^ child of the cluster i, M_i_ and S_i_ are, respectively, the estimated mean and standard deviation of the cluster i, which are the maximum of the mean and standard deviation predicted by the LOESS regression from the abundance of the parent sequence in cluster i and the mean and standard deviation estimated directly from the children abundance of the cluster i (Eqs 4 and 5). Taking the maximum of the mean and standard deviation enables us to be more conservative especially for low abundance sequences where data becomes sparse and the LOESS fit is less reliable. P-values generated by the “pnorm” test were adjusted for multiple hypothesis testing using the Benjamini-Hochberg Procedure. Corrected p-values less than 0.05 were considered significant, rejecting the null hypothesis that the variant child is a sequence error.

### Comparison to DADA2

We compared the performance of HashSeq to the performance of the DADA2 pipeline^5^. For this purpose, reads for each dataset were trimmed to the same length as discussed above. Trimming was performed with the function “filterAndTrim” in DADA2 with default parameters. Inference of sequence variants were performed as described in https://benjjneb.github.io/dada2/bigdata.html with default parameters. Filtering and inference were preformed using separate scripts in order to better compare the run time and memory usage between the DADA2 algorithm and HashSeq.

In order to compare the DADA2 and our algorithm, we used “blastn” to map sequence variants inferred from both algorithms to the SILVA132 database. Percent identity was calculated as:

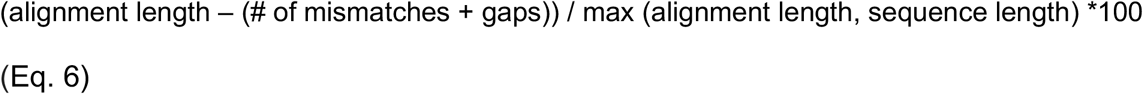

Where “sequence length” is the known length of the query sequence and all other parameters were reported by BLAST. This formula penalizes both mismatches and gaps in either sequence or in the alignment. For each sequence variant, the first hit with the highest bit score with the database was selected.

Finally, we compared DADA2 and our algorithm in terms of associations between the inferred sequence variants and the variable of interest in the metadata. For this purpose, we preformed Random Forest classification for each algorithm and dataset with four cross-validations and ten repeats using RandomForestClassifier with 100 decision trees and RepeatedKFold methods from Scikit-learn library in python 3.8.1.

## Data availability

Our pipeline is written in Java (JDK 1.8) and R (4.0.2) but can be installed as an R package and run from an R environment. Source code with instructions for installing HashSeq package can be found at https://github.com/FarnazFouladi/HashSeq. All codes and figures for the analyses of this manuscript can be found at https://github.com/FarnazFouladi/HashSeq_Manuscript.

## Supplementary Figure and Table Legends

**Supplementary Figure 1. The simple one-parameter Poisson model underestimates the abundance of sequence errors**. Plots show the relationship between the abundance of parent sequences and the abundance of one-mismatch children sequences on the log_10_ scale. The Poisson test with error rates estimated from the Poisson model (the best fit for each dataset; green lines in Figure 1) was used to test the null hypothesis that a one-mismatch child can be explained by sequencing error. Red and black dots indicate significant and insignificant p-values, respectively, at FDR 5%.

**Supplementary Figure 2. The fraction of all possible unique one-mismatches for a sequence can be well fit to a Poisson model as well as simulated data**. Black circles in the plot show the relationship between the abundance of parents on a log_10_ scale and the fraction of all possible unique one-mismatches that are observed for each parent sequence in the China dataset. The red line shows the fraction of all possible one-mismatches predicted by a one-parameter Poisson model that includes an error rate of 0.00015. The blue line shows the fraction of all possible one-mismatches that are simulated from Java code (see methods) with an error rate 0.00015 for sequences with different sequence depths.

**Supplementary Table 1. Datasets used in this study**. The table includes the project numbers associated with each dataset and the information regarding sequencing, including the sequencing instrument, the variable region in the 16S rRNA gene, and primers where available.

## Notes

https://github.com/FarnazFouladi/HashSeq

https://github.com/FarnazFouladi/HashSeq_Manuscript

### Competing Interest Statement

The authors have declared no competing interest.

