## Supplementary figures and images for "HashSeq: A Simple, Scalable, and Conservative *De Novo* Variant Caller for 16S rRNA Gene Datasets"

### Supplementary Figure 1

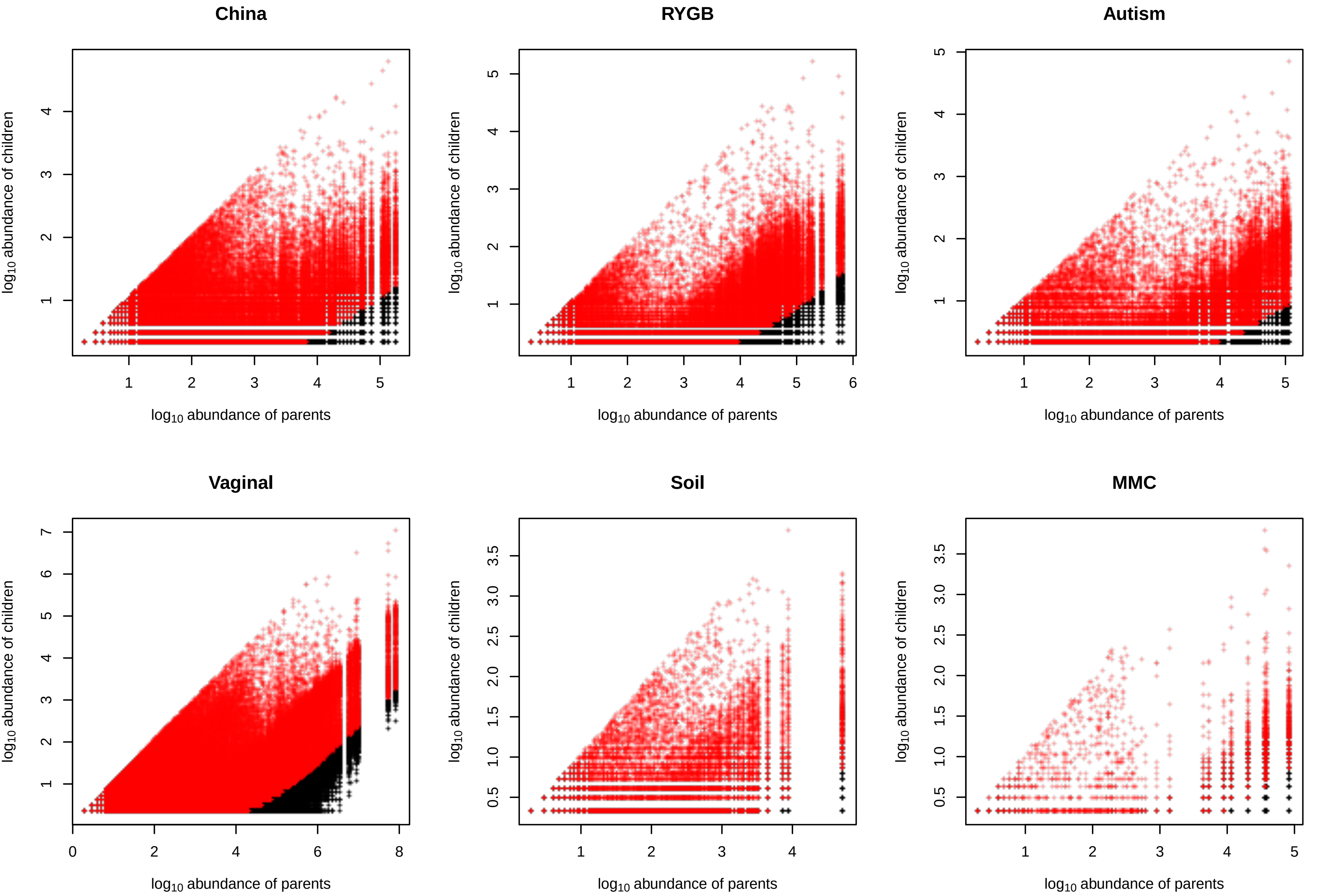

### Supplementary Figure 2

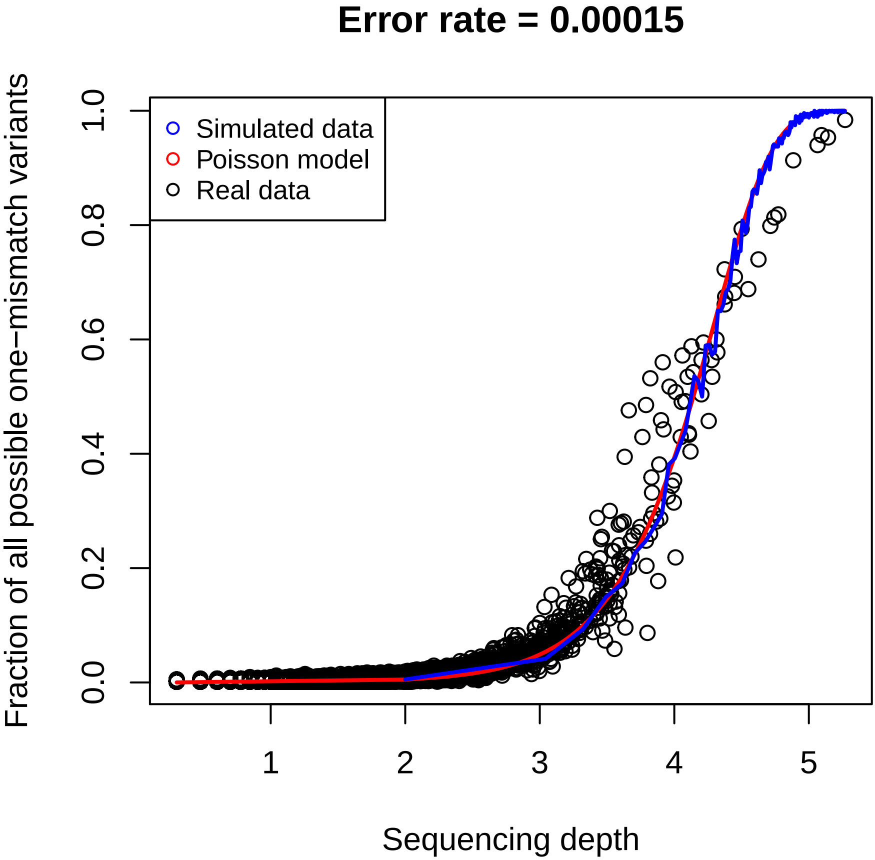
