## Supplementary Table 1 for "HashSeq: A Simple, Scalable, and Conservative *De Novo* Variant Caller for 16S rRNA Gene Datasets"

| Datasets | Project# | | Region | Forward primer | Reverse primer |
|---|---|---|---|---|---|
| autism | PRJNA533120 | MiSeq | V4 | GTGCCAGCMGCCGCGGTAA | GGACTACHVGGGTWTCTAAT |
| RYGB | SRP113514 | MiSeq | V4 | TCGTCGGCAGCCAGTGATGTGTATAAGAGACAGGTGCCAGCMGCCGCGGTAA | GTCTCGTGGGCTCGGAGATGTGTATAAGAGACAGGGACTACHVGGGTWTCTAAT |
| China | PRJNA349463 | MiSeq | V4 | Not available | Not available |
| Soil | PRJEB14409 | MiSeq | V3-V4 | CCTACGGGNGGCWGCAG | GACTACHVGGGTATCTAATCC |
| vaginal | SRP115697 | HiSeq 2500 | V4 | Not available | Not available |
| MMC | PRJEB24409 | MiSeq | V4-V5 | CAGCMGCCGCGGTAATACG | Not available |
